# *In vivo* analysis of *Trypanosoma cruzi* persistence foci at single cell resolution

**DOI:** 10.1101/2020.05.13.092551

**Authors:** Alexander I. Ward, Michael D. Lewis, Archie Khan, Conor J. McCann, Amanda F. Francisco, Shiromani Jayawardhana, Martin C. Taylor, John M. Kelly

**Author notes:** Address correspondence to John M. Kelly.

## Abstract

Infections with *Trypanosoma cruzi* are usually life-long despite generating a strong adaptive immune response. Identifying the sites of parasite persistence is therefore crucial to understand how *T. cruzi* avoids immune-mediated destruction. However, this is a major technical challenge because the parasite burden during chronic infections is extremely low. Here, we describe an integrated approach involving comprehensive tissue processing, *ex vivo* imaging, and confocal microscopy, which has allowed us to visualise infected host cells in murine tissue, with exquisite sensitivity. Using bioluminescence-guided tissue sampling, with a detection level of <20 parasites, we show that in the colon, smooth muscle myocytes in the circular muscle layer are the most common infected host cell type. Typically, during chronic infections, the entire colon of a mouse contains only a few hundred parasites, often concentrated in a small number of cells containing >200 parasites, that we term mega-nests. In contrast, during the acute stage, when the total parasite burden is considerably higher and many cells are infected, nests containing >50 parasites are rarely found. In C3H/HeN mice, but not BALB/c, we identified skeletal muscle as a major site of persistence during the chronic stage, with most parasites found in large mega-nests within the muscle fibres. Finally, we report that parasites are also frequently found in the skin during chronic murine infections, often in multiple infection foci. In addition to being a site of parasite persistence, this anatomical reservoir could play an important role in insect-mediated transmission, and have implications for drug development.

**IMPORTANCE:** *Trypanosoma cruzi* causes Chagas disease, the most important parasitic infection in Latin America. Major pathologies include severe damage to the heart and digestive tract, although symptoms do not usually appear until decades after infection. Research has been hampered by the complex nature of the disease and technical difficulties in locating the extremely low number of parasites. Here, using highly sensitive imaging technology, we reveal the sites of parasite persistence in experimental mice at single-cell resolution. We show that parasites are frequently located in smooth muscle cells in the circular muscle layer of the colon, and that skeletal muscle cells and the skin can also be important reservoirs. This information provides a framework for investigating how the parasite is able to survive as a life-long infection, despite a vigorous immune response. It also informs drug-development strategies by identifying tissue sites that must be accessed to achieve a curative outcome.

## INTRODUCTION

The intracellular protozoan parasite *Trypanosoma cruzi* is the etiological agent of Chagas disease, and can infect a wide variety of mammalian hosts. Transmission to humans is mainly via the hematophagous triatomine insect vector, which deposits infected faeces on the skin after a blood-meal, with the parasite then introduced through the bite wound or mucous membranes. Oral, congenital and blood transfusion are other important transmission routes. 6-7 million people in Latin America are infected with *T. cruzi* (1), and as a result of migration, there are now hundreds of thousands of infected individuals in non-endemic regions, particularly the USA and Europe (2, 3).

In humans, infection normally results in mild symptoms, which can include fever and muscle pain, although in children the outcome can be more serious. Within 6 weeks, this acute phase is usually resolved by a vigorous CD8+ T cell response (4, 5), and in most cases, the infection progresses to a life-long asymptomatic chronic stage, where the parasite burden is extremely low and no apparent pathology is observed. However, in ∼30% of individuals, the infection manifests as a symptomatic chronic condition, although this can take many years to develop. The associated cardiac dysfunction, including dilated cardiomyopathy and heart failure, is a major cause of morbidity and mortality (6, 7). In addition, ∼10% of those infected display digestive pathologies, such as megacolon and megaoesophagus, which on occasions can occur in parallel with cardiac disease. There is no vaccine against *T. cruzi* infection, and the current frontline drugs, benznidazole and nifurtimox, have limited efficacy, require long treatment regimens, and can result in severe side effects (8, 9). The global effort to discover new drugs for Chagas disease involves not-for-profit drug development consortia, as well as the academic and commercial sectors (10, 11). Progress would benefit considerably from a better understanding of parasite biology and disease pathogenesis.

One of the major challenges in Chagas disease research is to determine how *T. cruzi* survives as a life-long infection, despite eliciting a vigorous immune response which is able to reduce the parasite burden by >99%. Exhaustion of the parasite-specific CD8+ T cell response does not appear to be the reason (12). Alternative explanations include the possibility that *T. cruzi* is able to persist in immune-tolerant tissue sites (13), and the potential for the parasite to assume a non-dividing dormant form that does not trigger an overt immune response (14). Attempts to investigate these issues in humans have been limited by the long-term and complex nature of the disease, and by difficulties in monitoring tissue infection dynamics during the chronic stage. By necessity, most information on the sites of parasite location in humans has come from autopsy and transplant studies (15), and the significance of these data to patients in the asymptomatic chronic stage is unclear. Bioluminescence imaging of animal models has therefore been adapted as an approach to explore aspects of host:parasite interaction, disease pathology and drug-development (16-18). Our previous work has exploited highly sensitive *in vivo* imaging to monitor mice infected with bioluminescent *T. cruzi* that express a red-shifted luciferase (19-21). These experiments have shown that mice are useful predictive models for human infections in terms of infection dynamics (21, 22), drug-sensitivity (23) and the spectrum of cardiac pathology (24). We have also demonstrated that *T. cruzi* infection is pan-tropic during the acute stage, and that the adaptive immune response results in a 100 to 1000-fold reduction in the whole animal parasite burden as infections transition to the chronic phase, a process initiated 2-3 weeks post infection. The gastrointestinal (GI) tract, particularly the colon and/or stomach, was found to be a major site of parasite persistence during chronic stage infections, but it has not so far been possible to identify the infected host cell types in these complex tissues. The immune-mediated restriction to the GI tract was not absolute, with both host and parasite genetics impacting on the extent to which the infection could disseminate to a range of other organs and tissues (22). The severity of chronic cardiac pathology in different mouse strains was associated with the ability of parasites to spread beyond the permissive niche provided by the GI tract, and with the incidence of cardiac infection. This led us to propose a model in which the development of chagasic cardiac pathology, was linked with the frequency of the localised inflammatory immune responses stimulated by periodic trafficking of parasites into the heart (13).

More detailed information on the precise sites of parasite survival during chronic infections will provide new insights into disease pathogenesis, and aid the design of both immunotherapeutic and chemotherapeutic strategies. The scarcity of parasites during the chronic stage has made addressing this issue a major challenge, with PCR-based approaches being both uninformative with respect to host cell types, and unreliable because of the highly focal and dynamic character of infections (20, 23). To resolve this, we constructed *T. cruzi* reporter strains engineered to express a fusion protein that was both bioluminescent and fluorescent (25). This allowed individual infected host cells to be visualised routinely within chronically infected mouse tissue. The bioluminescent component facilitates the localisation of infection foci within *ex vivo* tissue samples, and fluorescence then enables histological sections to be rapidly scanned to identify infected cells (26). The utility of this approach has been further extended by using EdU labelling and TUNEL assays to explore the replicative status of parasites *in situ*.

Here, we describe how these enhanced imaging procedures, coupled with modifications to tissue processing, have allowed us to identify the sites of parasite persistence during chronic murine infections. We reveal that the circular muscle layer is the major reservoir of infection in the colon, that skeletal muscle can be an important site of persistence, although this phenomenon appears to be strain-specific, and that the skin can harbour multiple infection foci.

## RESULTS

### Locating the sites of *T. cruzi* persistence within the external wall of the colon during chronic murine infections

In multiple murine models, with a variety of parasite strains, bioluminescence imaging has revealed that the GI tract, particularly the large intestine and stomach, is a major site of parasite persistence during chronic *T. cruzi* infection (20, 22). However, our understanding of how this impacts on pathogenesis has been complicated by the difficultly in precisely locating, and then visualising, parasite infected cells. To resolve these technical issues, we infected mice with the *T. cruzi* CL-Luc::Neon line that constitutively expresses a reporter fusion protein that is both bioluminescent and fluorescent (25), and adapted our dissection procedures to allow a more detailed assessment of parasite location (Materials and Methods). At various periods post-infection, the colon of each mouse was removed, pinned luminal side up, and peeled into two distinct sections (Fig. 1a and b) - the mucosal layers consisting of (i) thick mucosal, muscularis mucosa, and submucosal tissue, and (ii) the muscular coat, including the longitudinal and circular smooth muscle layers, the enteric neuronal network, at the level of the myenteric plexus, intramuscular neurons and extrinsic nerve fibres. The resulting external gut wall mount is thin enough, and sufficiently robust, to allow the full length of the colon to be viewed in its entirety at a 3-dimensional level by confocal laser scanning microscopy. Using this approach, each bioluminescent focus in live peeled tissue from chronically infected mice could be correlated with fluorescent parasites in individual infected host cells (Fig. 1c and d). The resulting images revealed that the limit of detection achievable by bioluminescence imaging is less than 20 parasites. This level of sensitivity, in an *ex vivo* context, confirms the utility of this model for studies on infection dynamics (22), and drug and vaccine efficacy (24, 27, 28). In infected host cells, the number of parasites could be determined with precision using full-thickness serial Z-stacking (Fig. 1e, Fig. S1). This allowed us to establish that the total number of parasites persisting in the external colonic wall (tunica muscularis) of a chronically infected mouse is typically in the range of a few hundred (697 ± 217, n=16), although this number can be higher if the tissue contains one or more “mega-nests” (Fig. 1c, highlighted in yellow, as example).

**FIG 1.**
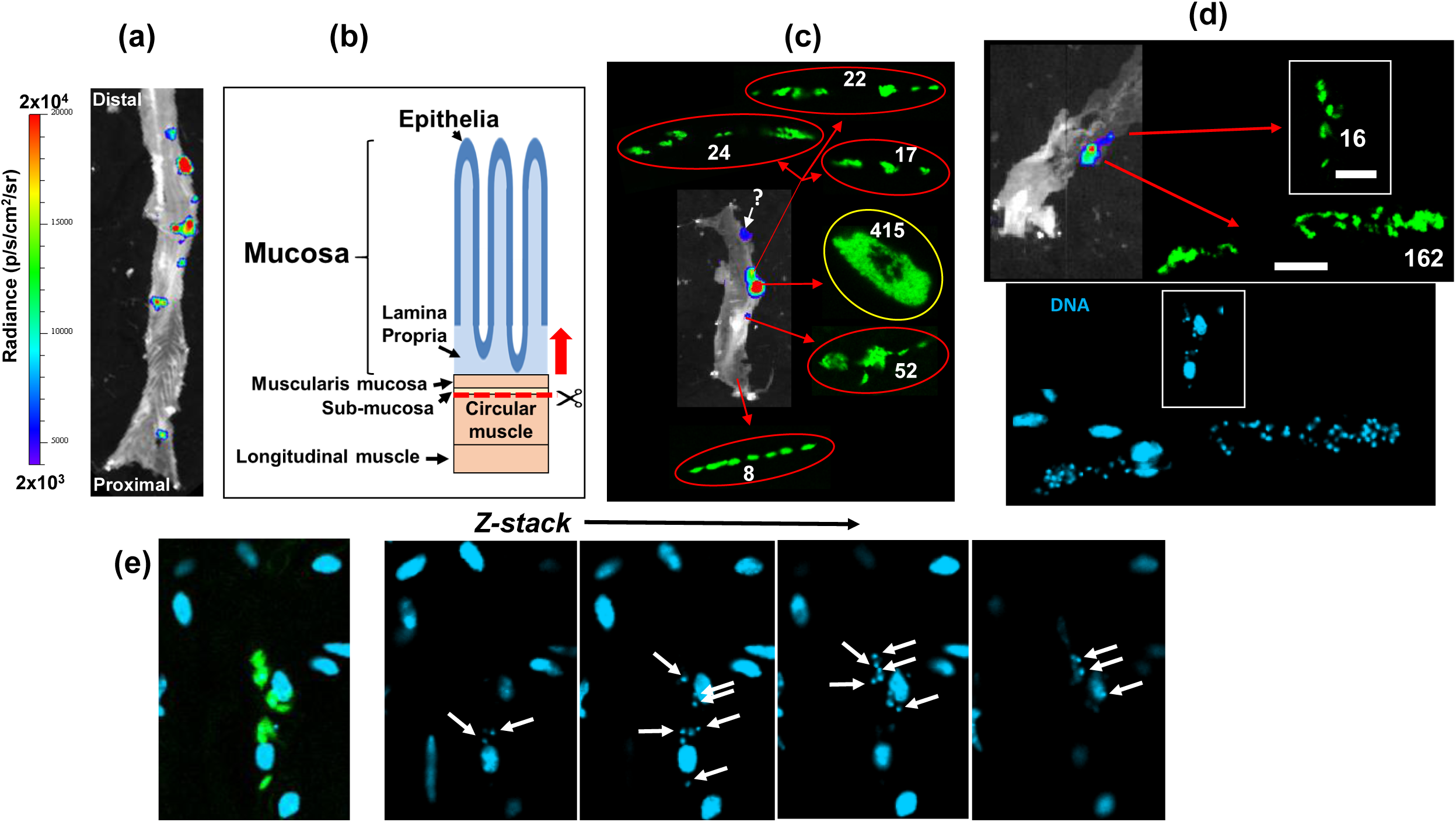
The limit of detection by *ex vivo* bioluminescence imaging of the murine colon is less than 20 parasites. (a) *Ex vivo* bioluminescence imaging of a section of the colon from a C3H/HeN mouse chronically infected (155 days post-infection) with *T. cruzi* CL-Luc::Neon (25), pinned luminal side up. The bioluminescence signal is on a linear scale pseudocolour heat map (same for all bioluminescence images in this figure). (b) Schematic showing the distinct layers of the GI tract (see also Fig. 3a). The dashed red line indicates the position above which tissue can be peeled off to leave the external colonic wall layers. (c) Bioluminescence image of a colonic wall section after peeling. The insets show the fluorescent parasites (green) detected after exhaustive 3-dimensional imaging of the tissue section (Materials and Methods), and the numbers detected. Parasites corresponding to one bioluminescent focus (marked by ‘?’) could not subsequently be found, due to technical issues. (d) Upper image; an external colonic wall layer from a separate mouse showing the correlation of bioluminescence imaging and fluorescence (green), including an infection focus with 16 parasites (left) (white scale bars=20 μm). Lower image; staining with DAPI identifies the location of parasite (small) and host cell (large) DNA. (e) Determination of parasite number. Serial Z-sections of the external colonic wall tissue containing the parasite nest shown in (d) indicate how 3-dimensional imaging can be used to calculate the number of parasites on the basis of DNA staining. See Fig. S1 for more detail.

When we compared parasite distribution in the external gut wall during acute and chronic murine infections, the most striking difference was the presence in the chronic stage of some host cells that were infected with >200 parasites (Fig. 2). The existence of these mega-nests resulted in a significant alteration in parasite number distribution at the level of single infected host cells (Fig. 1, Fig. 2b-d). In acute infections, parasites were spread between many more host cells, with the average parasite content per cell remaining relatively low (Mouse (M)1=6.5, M2=6.7, M3=6.5, M4=4.6, M5=19.7, mean=9.4, 1.15<µ<16.46, 95% confidence interval) (Fig. 2a, c and d). In the chronic stage, the situation was different. Of the total parasite number in the smooth muscle, more than half were present in mega-nests of >200, although most infected cells (>90%) contained fewer than 50 parasites. Nest size could extend to >1000 parasites. The number per infected cell was determined by Z-stacking, which could be done with accuracy, even at this level of parasite burden (for details, see Fig. S1). In the chronic stage, fully developed trypomastigotes were not observed in any of the infected cells examined during this study. In contrast, fully developed flagellated trypomastigotes were routinely observed in nests during the acute stage (Fig. 2e, as example). We did not find a single mega-nest in external colonic wall tissue derived from any animal during an acute stage infection, with 63 parasites being the maximum.

**FIG 2.**
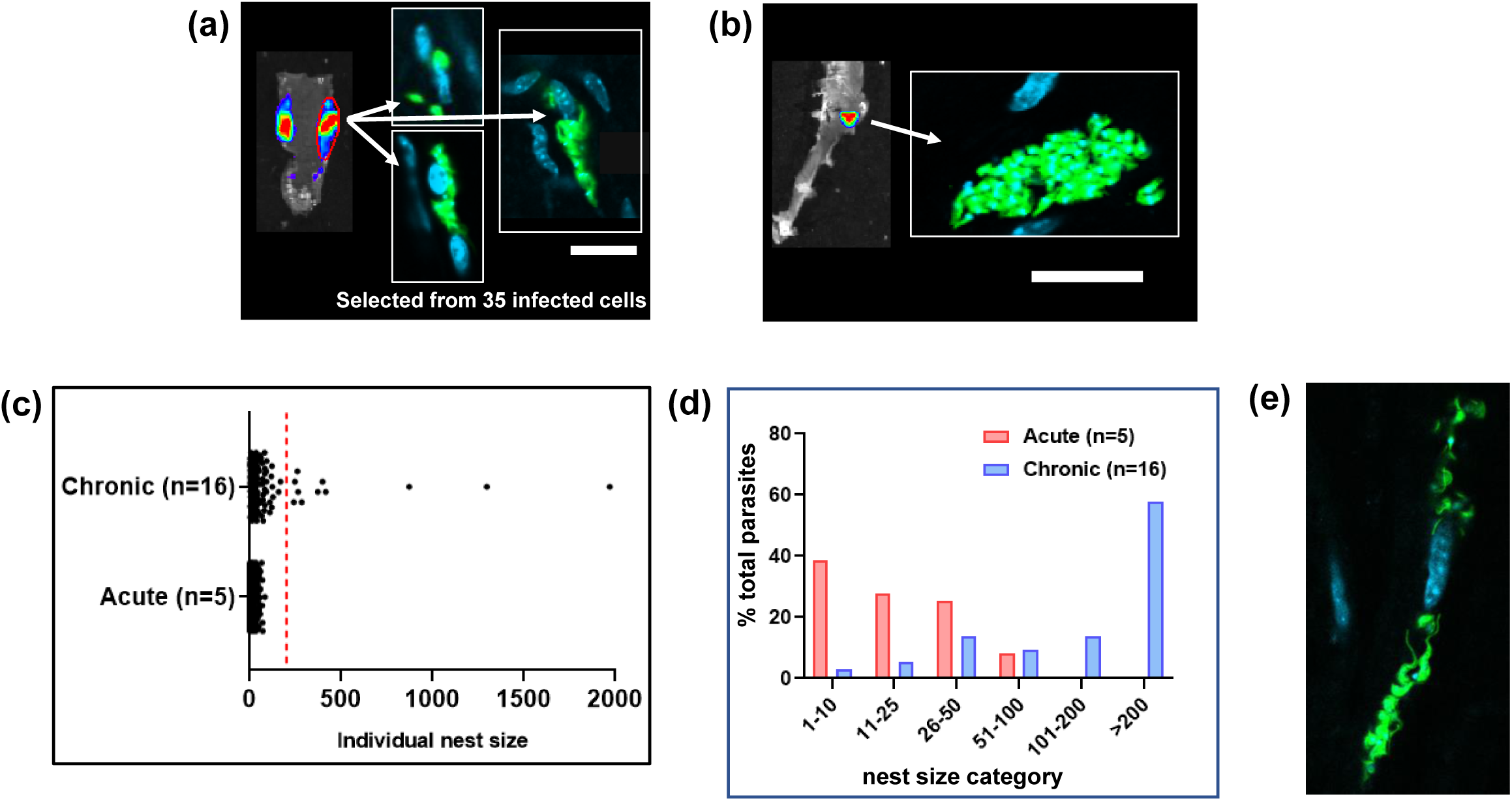
In the external colonic wall of chronic stage mice, cells infected with more than 200 parasites contain much of the *T. cruzi* population. (a) Bioluminescence imaging of peeled colon isolated from a C3H/HeN mouse 15 days post-infection (acute stage). After mounting, the region of interest (ROI) encompassed by the red line was exhaustively searched by confocal microscopy. 35 infected cells were found within the ROI, 3 of which are shown. Parasites, green. (b) Using the same approach, the external colonic wall from a chronically infected mouse (183 days post-infection) was assessed. The bioluminescent focus corresponded to a single highly infected host cell. White scale bars=20 μm. (c) Pooled data from *T. cruzi* infected cells in peeled colonic wall tissue muscle, isolated from 5 acutely and 16 chronically infected mice. Tissue was examined and the number of parasites per host cell established after the use of Z-stacking to provide a 3-dimensional image (Fig. S1). Each dot represents a single infected cell (acute stage, n=1198; chronic stage, n=140). (d) The same data set expressed as the % of the total parasites detected in the colons of mice in the acute (n=5) and chronic (n=16) stage of infection, by nest size category. (e) An infected cell in the colon of a mouse in the acute stage (15 days post-infection) of infection in which the parasites have differentiated to flagellated trypomastigotes.

To more accurately determine the specific location of parasites within the colon of chronically infected mice, we made histological sections of paraffin embedded whole colon tissue derived from both C3H/HeN and BALB/c mice infected with the CL Brener dual reporter strain. Using bioluminescence-guided sampling and confocal imaging, we exhaustively searched the tissue sections for fluorescent parasites (>100 sections per mouse). Bioluminescent foci could be well correlated with individual infected host cells, or small numbers of infected cells in close proximity (Fig. 3b, Fig. S2). Infected cells were most commonly located in the circular muscle layer, and only infrequently in the longitudinal muscle, or, despite its relatively larger size/volume, the mucosal layer (Fig. 2, Fig. 3b and c, Fig. S2). No infections of the columnar epithelial cells in the mucosal layer were detected in any mouse. We therefore conclude that in the colon, smooth muscle tissue is the major, although not the exclusive site of parasite persistence during chronic infection. Consistent with the whole mount imaging results (Fig. 2c), there was high variability in the number of *T. cruzi* per infected cell in the colonic tissue, ranging from single parasites to nests of >200, but no obvious correlation between the parasite burden per cell and the location of the infected cells within the various tissue layers. In the whole tissue mounts, based on the bioluminescence profile, there was a tendency for the proximal region of the colon to be more highly infected than the mid and distal regions, although this did not reach statistical significance (Wilcoxon rank sum test) (Fig. 4a).

**FIG 3.**
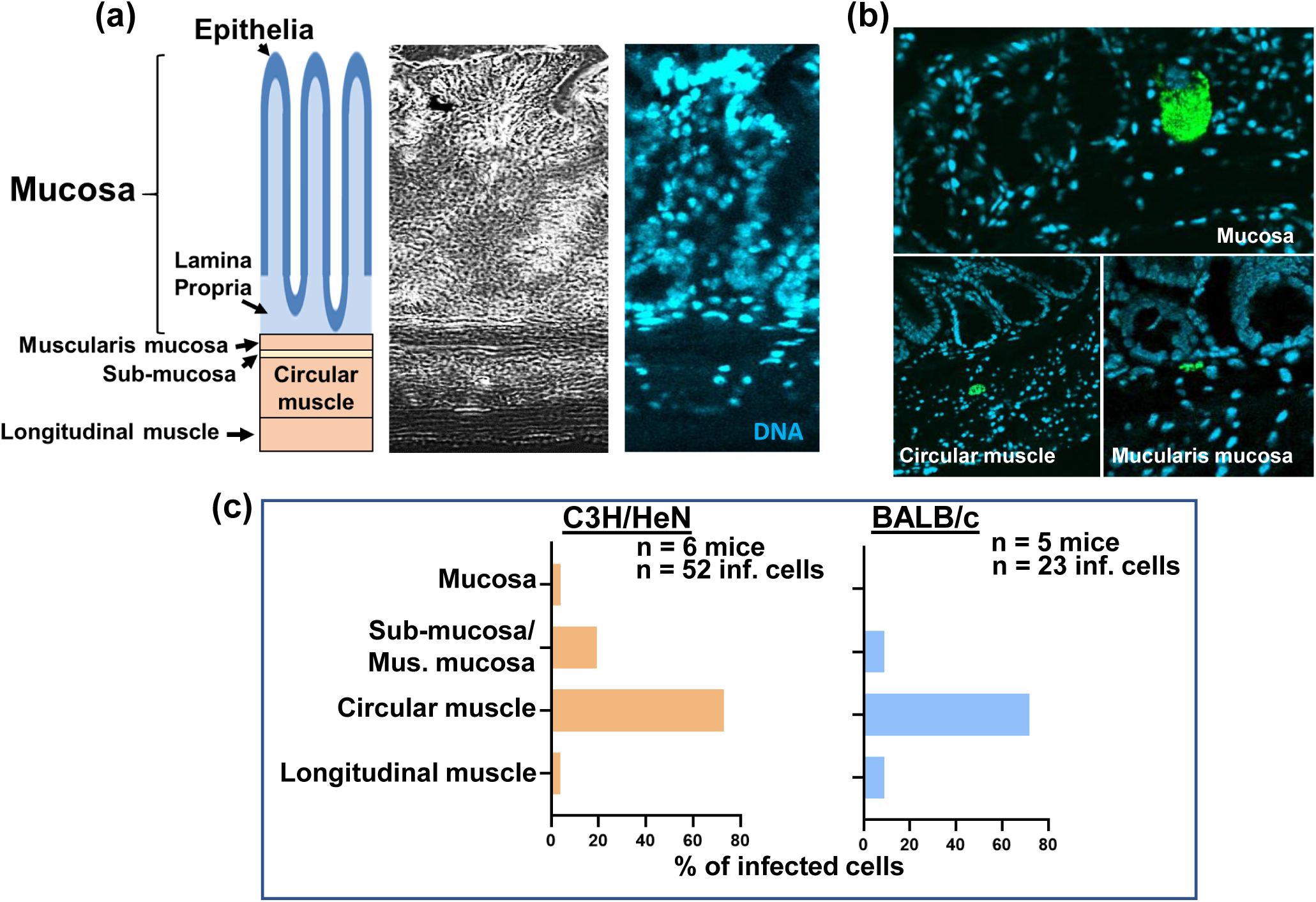
The majority of parasites in the colon of a chronically infected mouse are located in the circular muscle section. (a) Depiction of the layers of the GI tract, correlated with the phase (left) and DNA stained (DAPI) (right) images of the same tissue section. (b) Examples of host cells infected with fluorescent parasites (green) detected in different layers of the GI tract (see also Fig. S2). Infection foci were located by confocal imaging of fixed histological sections. (c) Summary of parasite location data obtained from chronically infected C3H/HeN and BALB/c mice.

**FIG 4.**
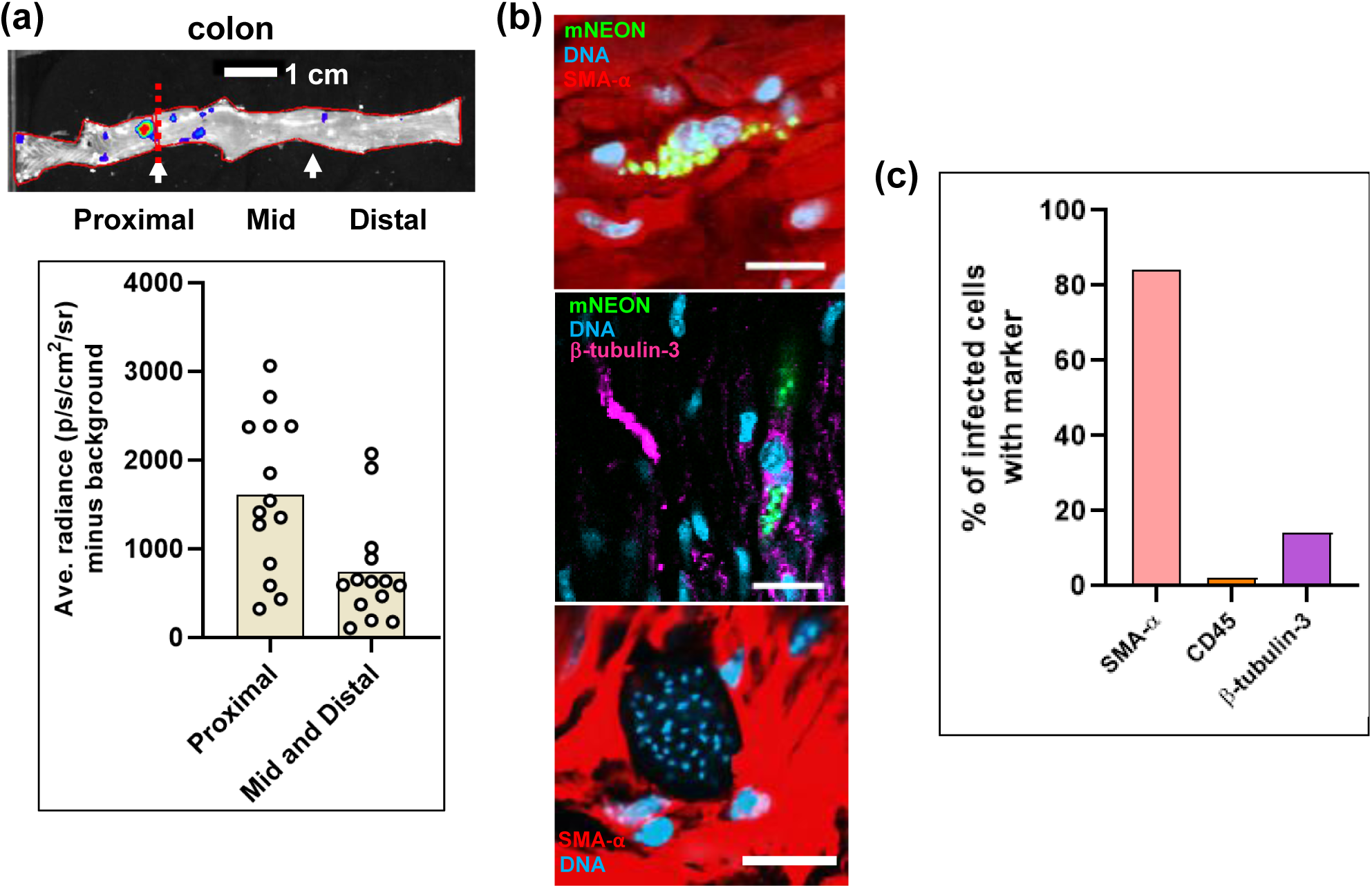
Smooth muscle cells are the predominant infected cell type in the GI tract of chronically infected mice. (a) Bioluminescence image of the large intestine of a chronically infected C3H/HeN mouse indicating the proximal, mid and distal regions, defined as the first, second and third segments measured using image J software. Data were analysed as described (Materials and Methods) (n=14) and are presented in the bar chart as the average radiance (p/s/cm^2^/sr) minus the background. (b) Illustrative images taken with the mounted external colonic wall section, following staining with cell type specific antibodies (Materials and Methods). Upper, infected smooth muscle cell. Middle, infected neuronal cell. Lower, a large parasite nest, refractive to staining with any of the 3 markers. (c) Bar chart summarising distribution of infection by host cell type. External colonic wall sections were single-stained with cell type specific antibodies. For smooth muscle (SMA-α; n=4 mice, 24 infected cells, 20+ve); for neuronal cells (β-tubulin-3; n=3 mice, 14 infected cells, 2+ve); for immune cells (CD45, n=8 mice, 61 infected cells, 1+ve).

To identify the major cell type(s) which act as parasite hosts during chronic infections of the GI tract, we single-stained whole mounted external colonic wall sections with specific antibodies against SMA-α (smooth muscle actin-α), β-tubulin-3 (a marker for neurons), and CD45 (a broad range marker of all nucleated hematopoietic cells) (Materials and Methods). These experiments showed that smooth muscle myocytes were the predominant host cell type (Fig. 4b and c), with a minority of infected cells stained with the neuronal or leukocyte marker. Interestingly, mega-nests, cells infected with >200 parasites, were refractive to staining with any of the three markers (Fig. 4b).

### Assessing skeletal muscle and the skin as sites of parasite persistence during chronic stage murine infections

For this study, BALB/c and C3H/HeN mice were chronically infected with *T. cruzi* CL-Luc::Neon (25), and the dissection procedures used for *ex vivo* imaging (Fig. 5a) were further modified to extend the range of tissue sites that could be assessed (Materials and Methods). Total removal of the skin and fur from the carcass allowed the whole of the skeletal muscle system to be exposed and imaged (Fig. 5b and d). The skin could also be placed fur side down and imaged in its entirety after the removal of adipose tissue. All adipose tissue harvested during the dissection process was combined to be imaged separately.

**FIG 5.**
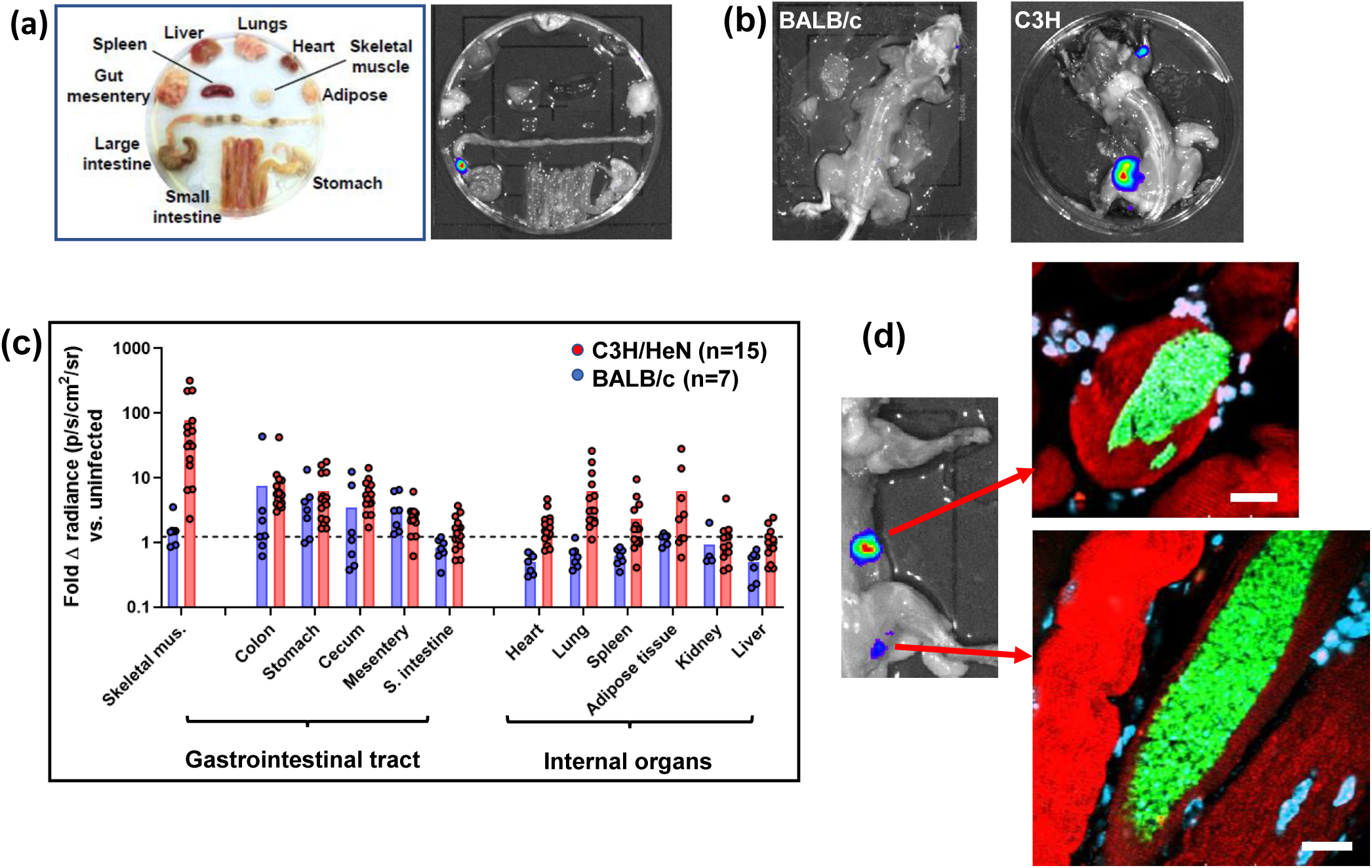
Skeletal muscle is a major site of parasite persistence during chronic *T. cruzi* infections in C3H/HeN mice, but not BALB/c. (a) *Ex vivo* imaging of organs and tissues from a BALB/c mouse chronically infected with bioluminescent *T. cruzi* CL Brener. (b) Dorsal bioluminescence imaging of chronically infected BALB/c and C3H/HeN mice following removal of internal organs, fur, skin and major adipose depots (Material and Methods). (c) Fold change in radiance (photons/s/cm^2^/sr) established by *ex vivo* bioluminescence imaging of internal tissues and organs and skeletal muscle as imaged in (a) and (b). Dashed line indicates the detection threshold equal to the mean +2SDs of the bioluminescence background derived from corresponding empty regions of interest obtained in tissue from age-matched uninfected mice. For technical reasons, on a small number of occasions, data could not be acquired for tissue samples from some mice (eg adipose tissue). (d) Bioluminescent foci from skeletal muscle were excised, histological sections prepared, and then scanned by confocal microscopy (Materials and Methods). Sections were stained with specific markers for muscle (actin-α, red) and DNA (DAPI; blue/turquoise). Parasites can be identified by green fluorescence. White scale bars=20 μm.

Each C3H/HeN mouse registered a bioluminescence signal in the skeletal muscle during chronic stage infections (n=16) (Fig. 5c). It could be inferred from the bioluminescence intensity that the parasite burden in this strain was significantly higher in skeletal muscle than in other organs or tissues, including the GI tract and lungs (*p*-value <0.001, Wilcoxon signed rank test) (Fig. 5b and c). As previously reported (22), parasite burden and dissemination during chronic stage infections is more extensive in C3H/HeN mice than in other mouse models. In line with this, we did not routinely detect highly bioluminescent foci in the skeletal muscle of BALB/c mice (Figure 5b and c). In addition, the adipose samples of the BALB/c mice were consistently close to background levels, whereas with the C3H/HeN mice, more than half displayed a detectable signal (>2SDs above background radiance) (Fig. 5c). Following bioluminescence-guided excision (Fig. 5d), infected foci from C3H/HeN skeletal muscle were subjected to histological sectioning and examined by confocal microscopy, with parasites detected on the basis of green fluorescence. Consistent with the external colonic wall data (Fig. 1), strong bioluminescent foci corresponded with large mega-nests constituted by many hundreds of parasites (Fig. 5d). Co-staining of these skeletal muscle sections with anti-actin-α antibodies revealed that the mega-nests were internal to the muscle fibres. Therefore, skeletal muscle represents an important site of parasite persistence in chronically infected C3H/HeN mice, but not in BALB/c.

Previous studies have shown evidence of *T. cruzi* infection foci localised to skin samples (20, 22). However, the extent to which the skin could act as a potential reservoir site has not been evaluated systematically. To investigate this, we infected C3H/HeN and BALB/c mice with the bioluminescent *T. cruzi* lines JR (DTU I) or CLBR (DTU VI), and employed a modified dissection protocol that allowed near-complete skins from infected mice to be subjected to *ex vivo* imaging after removal of subcutaneous adipose tissue (Materials and Methods) (Fig. 6a). Depending on the infection model, between 80% and 90% of mice had at least one discernible focus of skin infection (Fig. 6b). For all four parasite:mouse strain combinations, we observed a wide range of skin parasitism patterns, as judged by both the number and the intensity of the bioluminescent foci (Fig. 6a and b). There was some evidence that C3H/HeN mice had more CL Brener skin parasites than BALB/c mice (Fig. 6b and c). Infections with the CL Brener strain produced more discrete foci and a higher inferred parasite load than JR infections, although some of this effect could be attributed to lower luciferase expression levels in the JR strain (22). Skin imaging was conducted after removal of subcutaneous adipose tissue by dissection, strongly suggesting that the majority of parasites were resident in the dermis. To visualise parasites at the cellular level, bioluminescence positive biopsies were processed for thin section fluorescence imaging from infections with parasites expressing the bioluminescent:fluorescent fusion protein (n∼300 sections from 5 mice). Visualisation of infected cells in the skin biopsies was more technically challenging than for other tissues. Only a single, apparently multinucleated infected cell was identified (Fig. 6d), containing approximately 30 parasites and located within 150 μm of the epidermis. Parasites in this anatomical location are likely to have a role in disease transmission.

**FIG 6.**
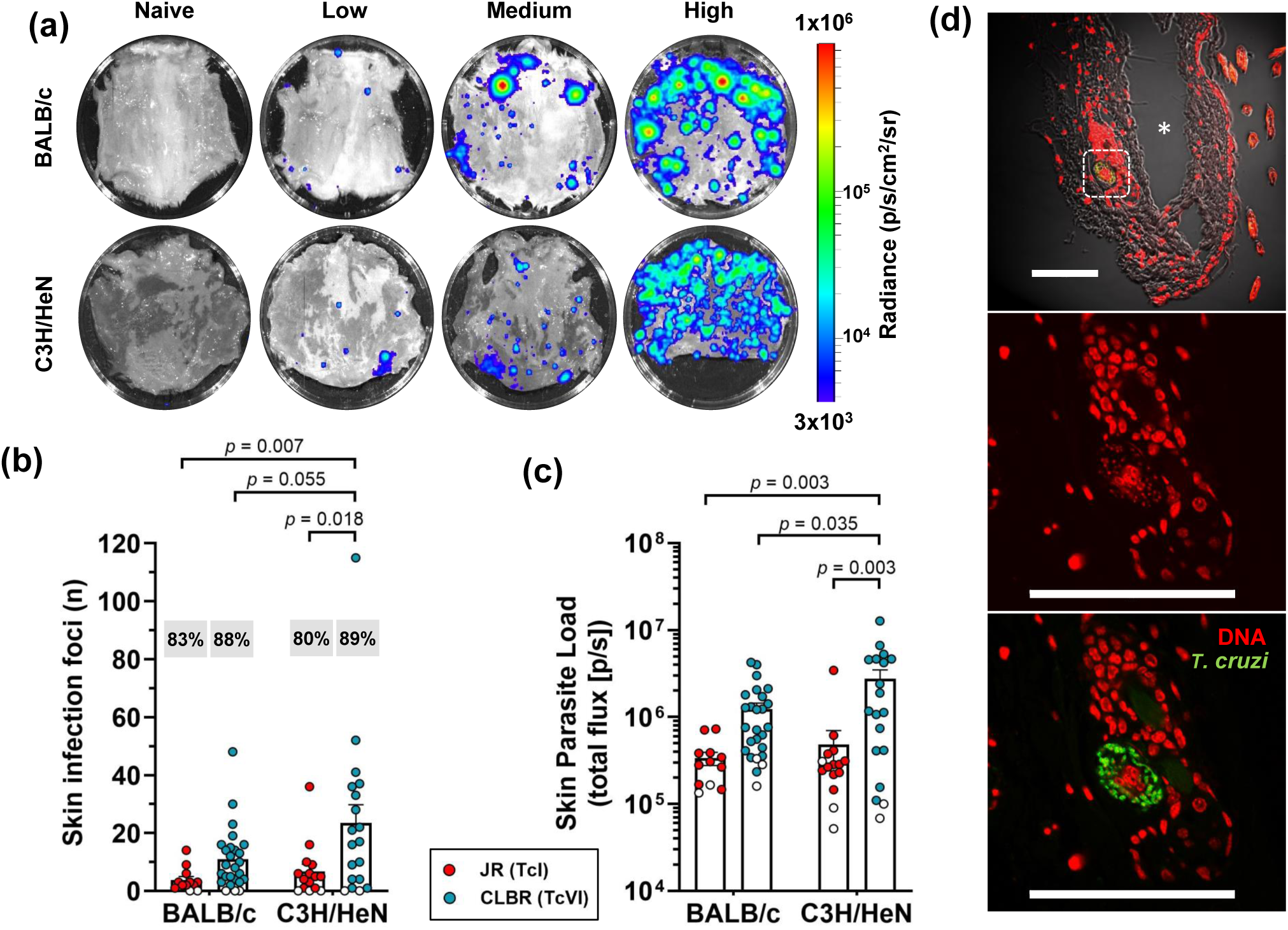
The skin is a major site of parasite persistence during chronic *T. cruzi* infections in mice. (a) *Ex vivo* bioluminescence imaging of skin (adipose tissue removed) from chronically infected BALB/c and C3H/HeN mice (>150 days post-infection) showing representative examples of low, medium and high parasite load. The bioluminescence signal is on a log10 scale pseudocolour heat map. (b and c) Quantification of the number of discrete infection foci (b) and the bioluminescence intensity for each skin (c). Data points are individual animals, with empty circles indicating skins having zero radiance above background. Mean values ±SEM are shown. Percentages in grey boxes (b) refer to the number of animals with at least one focus above the bioluminescence threshold. Infections with both *T. cruzi* CL Brener and JR bioluminescent strains were assessed (n=12-26 animals per combination, 3-4 independent experiments). Groups were compared by 2-way ANOVA. (d) Confocal micrographs showing fluorescent CL Brener parasites in an infected cell within the dermis of a BALB/c mouse 230 days post-infection (surface to the right). Upper image (200x). Asterisk indicates a gap resulting from a cutting artefact. Lower two images (630x) highlight the region in the white boxed area (above). White scale bars=100 μm.

## DISCUSSION

Chronic Chagas disease in humans is characterised by long-term parasite persistence at levels that are difficult to monitor with accuracy, even using highly sensitive PCR-based techniques. This has been a complicating factor in diagnosis, and in monitoring therapeutic cure during clinical trials (29, 30). Additionally, it has not been possible to identify the main tissues and/or organs that function as sites of parasite persistence in an immunological environment that otherwise tightly controls the infection. Information on the systemic parasite load and location throughout the infection would provide a better understanding of disease progression and the determinants of the wide spectrum of symptoms that are characteristic of this chronic condition. Experimental animal models have proved to be invaluable experimental tools for providing data in these areas, particularly in combination with genetically modified parasite reporter strains. These systems can provide real-time readouts on infection dynamics (20, 22, 31), insights into tissue tropism (26), and information on the influence of host and parasite genetics. The murine models used in the current study display a similar infection profile to that in humans, have proved to be predictive of drug efficacy, and display a spectrum of cardiac pathology that mirrors aspects of the human disease.

Here, we exploited parasites that express fusion proteins containing bioluminescent and fluorescent domains. Together with improved tissue preparation techniques, this has enabled us to achieve a limit of detection by *ex vivo* imaging that is less than 20 parasites (Fig. 2d and e). By facilitating the routine detection of parasites in their tissue context, at the level of individual host cells, these approaches have overcome a major barrier that has restricted progress in the investigation of chronic *T. cruzi* infections. Previous reports using bioluminescent parasites identified the GI tract as a major site of parasite persistence during the chronic stage (20, 22). However, these studies, which involved several mouse:parasite strain combinations, revealed few details on the nature of host cells, or on their precise location within tissue. In the colon, we have now shown that the circular smooth muscle coat is the predominant site of parasite persistence (Fig. 3) and that smooth muscle myocytes are the main infected host cell type. Enteric neurons can also be parasitized, but these infections are much less common (Fig. 4). The extent to which this apparent tropism is determined by a metabolic preference for the corresponding regions/cells, or by the immunological microenvironment is not known. Interestingly, external colonic wall resident CD45+ve hematopoietic cells were rarely infected (Fig. 4a), even though myeloid cells are well known targets during the acute stage infection in other sites such as the spleen or bone marrow. We also failed to find a single instance where parasites infected epithelial cells on the mucosal surface, suggesting that parasitized cells or trypomastigotes are unlikely to be shed into the lumen of the large intestine.

Experiments have shown that parasite survival in the colon during chronic infections reflects crucial differences between the immune environment of certain GI tract regions and systemic sites (22). Immunosuppression of infected mice leads to widespread parasite dissemination to other less permissive organs and tissues, including the heart. There is clearly a host genetic component to this immune restriction since the same parasite strains display a wider tissue distribution in C3H/HeN mice than in the BALB/c strain (Fig. 5c), a phenomenon which is associated with increased cardiac pathology (22). In the human population, this highlights that genetic diversity affecting the functioning of the immune system and its ability to restrict the tissue range of *T. cruzi* to reservoir sites could be a major determinant of Chagas disease pathology. Within C3H/HeN mice, skeletal muscle was also found to be an important site of persistence during the chronic stage, whereas in the BALB/c strain, parasites were far less evident in this location (Fig. 5c). Some *T. cruzi* strains have been reported to be myotropic in mice, with pathological outcomes that include paralyzing myositis and skeletal muscle vasculitis (32). Myocyte infections could also provide the parasite with access to myoglobin, a source of haem or iron that may contribute to a nutritional environment that is favourable for replication. The ability of high numbers of parasites to survive long-term in the skeletal muscle, compared to other sites, indicates that this tissue can function as a more immunologically permissive niche in the genetic background of the C3H/HeN mouse. Strikingly, myocytes in this tissue could contain several hundred parasites (Fig. 5d). We have previously suggested that the existence of large mega-nests such as these could have implications for drug efficacy (26), with parasites in the centre of the nest having reduced drug exposure compared to those on the periphery, possibly contributing to treatment failure. This form of “herd-protection” may not be captured in the type of high-throughput *in vitro* screening assays that are a common feature of the drug development process. It will also be interesting to explore whether some parasites within these mega-nests adopt a metabolically quiescent state, analogous to the dormant phenotype recently reported (14).

Our study has also demonstrated that the skin is another location where *T. cruzi* can be frequently detected during chronic infections. In both C3H/HeN and BALB/c mice, infection levels of >80% were observed, although there was considerable variability in the level of infectivity in terms of the number of bioluminescent foci and the total parasite load. The extent of this only became apparent when the entire skin of the mouse was examined by *ex vivo* imaging with the fur side down (Fig. 6), presumably because bioluminescent signals at the levels displayed by the majority of foci are masked by the fur when monitored by *in vivo* imaging. Skin-localised parasites are a common and well characterised feature of many *Leishmania* species infections. More recently, it has also been reported that *Trypanosoma brucei*, can also be detected in the skin of both mice and humans, and that these parasites could have important roles in persistence and transmission (33, 34). Until now, descriptions of cutaneous *T. cruzi* have been restricted to intermittent (chagoma and Romaña’s sign) or atypical manifestations of the acute stage (35), or to reactivation of chronic infections as a result of immunosuppression (36, 37). Parasites in the dermal layers (Fig. 6) have the potential to play a crucial role in transmission of Chagas disease since they would have ready access to the triatomine vector during a blood meal. It will also be important to determine whether these skin-resident parasites are persistent at this location, or whether they represent a transient population that is constantly re-seeded from other permissive niches, such as the GI tract (13). Resolution of this question will help to inform drug-design by revealing whether the ability to access parasites in the dermal layers has to be a pre-requisite property of novel therapeutics. In murine models of *T. brucei* infection, adipose tissue also forms an important parasite reservoir (38). This was not the case with chronic *T. cruzi* infections of BALB/c mice (Fig. 5c), where parasites were largely absent from these tissue sites. Bioluminescent foci were detected in the adipose tissues in approximately half of the chronically infected C3H/HeN mice. However, rather than a specific tropism, this may simply reflect the immunological context in C3H/HeN mice, which allows more extensive parasite distribution than in other mouse models (22).

In summary, we have provided new data on the sites of parasite persistence in murine models of chronic Chagas disease. This provides a framework for identifying the immunological parameters that determine whether a specific tissue site can act as a permissive niche, and for investigating the extent to which the parasite itself has a direct role in the process.

## MATERIALS AND METHODS

### Ethics

Animal work was carried out under UK Home Office project licenses (PPL 70/8207 and P9AEE04E4) and approved by the LSHTM Animal Welfare and Ethical Review Board. All procedures were conducted in accordance with the UK Animals (Scientific Procedures) Act 1986 (ASPA).

### Parasites, mice and infections

Two parasite reporter strains were used; the bioluminescent:fluorescent *T. cruzi* CL-Luc::Neon, a CL Brener clone (DTU VI) engineered to express a fusion protein containing red-shifted luciferase linked to the mNeonGreen chromophore (20, 25), and a JR clone (DTU I), which expresses red-shifted luciferase (19, 22). Parasites were grown as epimastigotes at 28°C in RPMI-1640 supplemented with 0.5% (w/v) tryptone, 20 mM HEPES pH 7.2, 30 mM haemin, 10% heat-inactivated fetal bovine serum, 2 mM sodium glutamate, 2 mM sodium pyruvate, 100 µg ml^−1^ streptomycin and 100 U ml^−1^ penicillin, with 150 μg ml^-1^ hygromycin (CL Brener) or 100 µg ml^−1^ G418 (JR) as selective drugs. Metacyclic trypomastigotes (MTs) were obtained by transfer to Graces-IH transformation medium (39). MTs were harvested after 4–7 days, when typically, 70–90% of parasites had differentiated. Tissue culture trypomastigotes were obtained from the infected MA104 kidney epithelial cell line grown at 37°C in 5% CO2 using Minimal Eagles medium supplemented with 10% heat-inactivated fetal bovine serum.

BALB/c and C3H/HeN mice were purchased from Charles River (UK), and CB17 SCID mice were bred in-house. Animals were maintained under specific pathogen-free conditions in individually ventilated cages. They experienced a 12 h light/dark cycle and had access to food and water *ad libitum*. Female mice aged 8-12 weeks were used for all infections. SCID mice were infected with 1×10^4^ *in vitro*-derived tissue culture trypomastigotes in 0.2 ml PBS via i.p. injection. All other mice were infected by i.p injection with 1×10^3^ bloodstream trypomastigotes derived from parasitemic SCID mouse blood. All infected SCID mice developed fulminant infections and were euthanized at, or before, humane end-points. At experimental end-points, mice were sacrificed by lethal injection with 0.1-0.2 ml Dolethal.

### *Ex vivo* bioluminescence imaging

For *ex vivo* imaging, mice were injected with 150 mg kg^−1^ d-luciferin i.p., then sacrificed by lethal i.p. injection 5 min later (20, 21). Mice were perfused with 10 ml 0.3 mg ml^−1^ d-luciferin in PBS via the heart. Organs and tissues were imaged in three stages using the IVIS Spectrum system (Caliper Life Science) and the LivingImage 4.7.2 software. Firstly, heart, lungs, spleen, liver, GI tract, GI mesenteric tissue, kidneys and all visceral adipose tissue were transferred to a Petri dish in a standardized arrangement, soaked in 0.3 mg ml^−1^ d-luciferin in PBS and then imaged using maximum detection settings (2 min exposure, large binning). Then, the skin was removed from the carcass and subcutaneous adipose tissue removed (40) and added to the visceral fat creating a whole ‘adipose tissue’ sample, which was imaged in the same way. The skin was placed fur down, soaked in 0.3 mg ml^-1^ d-luciferin and imaged under the same conditions as the internal organs. The skeletal muscle was placed dorsal side up and soaked in 0.3 mg ml^-1^ d-luciferin and imaged, as above.

To quantify infection intensities in *ex vivo* tissues, individual regions of interest (ROIs) were drawn to quantify bioluminescence expressed as radiance (photons/s/cm^2^/sr). Because different tissue types from uninfected control mice have different background radiances, we normalized the data from infected mice using matching tissues from uninfected controls (n=4) and used the fold-change in radiance, compared with these tissue-specific controls, as the final measure of *ex vivo* bioluminescence. Detection thresholds for *ex vivo* imaging were determined using the fold-change in radiance for ROIs in images obtained from infected mice compared with matching empty ROIs in images from uninfected control mice of comparable age.

In some experiments, the colon was removed after standard imaging, an incision was made down the line of mesentery attachment, and the tissue sample pinned out under a dissection microscope. Using ultrafine tweezers, large sections of the smooth muscular coat from the other layers were peeled off, whilst the tissue remained bathed in 0.3 mg ml^−1^ d-luciferin (40). After imaging, luciferin was removed by 2x washing with PBS. Tissue was fixed with 4% paraformaldehyde for 45 min, followed by 2x washes with PBS (40). External colonic wall tissue was then whole mounted in Vectashield and imaged as below.

Histological sections were created after bioluminescence-guided excision of infection foci from skeletal muscle and colon tissue (25, 26, 40). The biopsies were first incubated in 95% EtOH at 4°C overnight, and then washed in 100% EtOH (4×10 min), followed by xylene (2×12 min). The samples were embedded by placing in melted paraffin wax (2×12 min). The wax was allowed to set and the embedded pieces were sectioned into 5-20 µm histological sections using a microtome. The sections were melted and paraffin dissolved in xylene for 30 s, then washed in 95% EtOH (3×1 min), followed by 3 washes in PBS. Sections were then mounted in Vectashield and imaged using the Zeiss LSM880 confocal microscope. For precise counting of intracellular parasites, samples were imaged in 3-dimensions, with the appropriate scan zoom setting, and the files exported for analysis using image J software (see Fig. S1).

### Antibody staining

Deparaffinized sections were incubated at 4°C overnight in primary antibody diluted at 1:200 in PBS/0.5% fetal bovine serum. Antibodies against β-tubulin-3 (Biolegend, Cat#802001), CD45 (Tonbo Biosciences, Cat#70-0451), smooth muscle actin (Sigma, Cat#A2547), and skeletal muscle actin (Thermo Fisher, Cat#MA5-12542) were used to stain for neuronal, nucleated hematopoietic, smooth muscle and skeletal muscle cells, respectively. Secondary antibodies (all obtained from Thermo Fisher) diluted 1:500 in PBS were incubated on sections for 3 h at room temperature before mounting. Both primary and secondary antibodies were removed by 3×2 min washes in PBS. For staining of whole colon external wall sections, the tissue was submerged in the primary antibody dilution for 48 h at 4°C, and then submerged in the secondary dilution at room temperature for 3 h before 3×2min washes in PBS.

### Statistics

The Shapiro-Wilk test for normality, and the Wilcoxon rank sum non-parametric test were used to analyse the data presented in Fig. 4 and 5. Two-way ANOVA with Tukey’s post hoc correction testing was used for Fig. 6. All tests were performed in GraphPad Prism v.8.

## ACKNOWLEDGEMENTS

This work was supported by the following awards: UK Medical Research Council (MRC) Grants MR/T015969/1 to JMK and MR/R021430/1 to MDL, and MRC LID (DTP) Studentship MR/N013638/1 to AIW. The funders had no role in study design, data collection and interpretation, or the decision to submit the work for publication.

## SUPPLEMENTAL MATERIAL

**FIG S1** The determination of parasite numbers in highly infected host cells. (a) Bioluminescent image of a peeled large intestine from a C3H/HeN mouse chronically infected with *T. cruzi* CL-Luc::Neon. (b) The excised tissue was imaged by confocal microscopy (x100) to reveal a highly infected smooth muscle cell (parasites, green). (c) The same image showing DAPI staining (blue) to reveal DNA. (d) For Z-stack analysis, the image was split into grids using the ZEN software. Parasite load was determined from the number of discoid-shaped kinetoplasts. To facilitate accurate counting, the relatively faint staining of the nuclear genome can be reduced by adjusting the contrast. (e) A series of 4 representative Z-stacked images from a total of 13 slices taken to assess parasite number across the infected cell. A total of 60 parasites were assigned to this 3-dimensional grid. The total number of parasites in the nest was 1969.

**FIG S2** Location of parasites within the murine GI tract during chronic *T. cruzi* infection. C3H/HeN mice were chronically infected with *T. cruzi* CL-Luc::Neon and the colon was examined by confocal imaging of histological sections following DNA staining (DAPI) (Materials and Methods). Host cells infected with fluorescent parasites (green, indicated by white arrows) were detected in different layers of the GI tract, as indicated. White scale bars=20 μm.

